# Rise in frequency of *lasR* mutant *Pseudomonas aeruginosa* among keratitis isolates between 1993 and 2021

**DOI:** 10.1101/2023.08.22.554354

**Authors:** Robert M. Q. Shanks, Sarah Atta, Nicholas A. Stella, Chollapadi V. Sundar-Raj, John E. Romanowski, Arman S. Grewel, Hazel Q. Shanks, Sonya M. Mumper, Deepinder K. Dhaliwal, Alex Mammen, Jake D. Callaghan, Rachel C. Calvario, Eric G. Romanowski, Regis P. Kowalski, Michael E. Zegans, Vishal Jhanji

**Affiliations:** Charles T. Campbell Laboratory of Ophthalmic Microbiology, Department of Ophthalmology, University of Pittsburgh School of Medicine, Pittsburgh, Pennsylvania, USA; Department of Surgery, Geisel School of Medicine at Dartmouth, Hanover, NH, USA; Department of Microbiology and Immunology, Geisel School of Medicine at Dartmouth, Hanover, NH, USA

**Author notes:** Corresponding author: Robert M. Q. Shanks.

**Keywords:** keratitis, *Pseudomonas aeruginosa*, ocular infection, quorum sensing, transcription factor, protease

## Abstract

*Pseudomonas aeruginosa* causes severe vision threatening keratitis. LasR is a transcription factor that regulates virulence associated genes in response to the quorum sensing molecule N-3-oxo-dodecanoyl-L-homoserine lactone. *P. aeruginosa* isolates with *lasR* mutations are characterized by an iridescent high sheen phenotype caused by a build-up of 2-heptyl-4-quinolone. A previous study indicated a high proportion (22 out of 101) of *P. aeruginosa* keratitis isolates from India between 2010 and 2016 were sheen positive and had mutations in the *lasR* gene, and the sheen phenotype correlated with worse clinical outcomes for patients. In this study, a longitudinal collection of *P. aeruginosa* keratitis isolates from Eastern North America were screened for *lasR* mutations by the sheen phenotype and sequencing of the *lasR* gene. A significant increase in the frequency of isolates with the sheen positive phenotype was observed in isolates between 1993 and 2021. Extracellular protease activity was lower among the sheen positive isolates and a defined *lasR* mutant. Cloned *lasR* alleles from the sheen positive isolates were loss of function or dominant negative and differed in sequence from previously reported ocular *lasR* mutant alleles. Insertion elements were present in a subset of independent isolates and may represent an endemic source from some of the isolates. Retrospective analysis of patient information suggested significantly better visual outcomes for patients with infected by sheen positive isolates. Together, these results indicate an increasing trend towards *lasR* mutations among keratitis isolates at a tertiary eye care hospital in the United States.

## Introduction

The bacterium *Pseudomonas aeruginosa* is the most frequent cause of contact lens associated microbial keratitis and is of concern because keratitis caused by *P. aeruginosa* has rapid progression and poor clinical outcomes(1, 2). *P. aeruginosa* keratitis isolates resistant to fluoroquinolones and other antibiotics typically used to treat keratitis have been reported (3-6). Microbial keratitis caused by antibiotic resistant *P. aeruginosa* isolates correlates with worse clinical outcomes including an increase in corneal perforations from 12% of cases with normal *P. aeruginosa* to 52% with multidrug-resistant (MDR) *P. aeruginosa* (1, 7). Beyond resistance, *P. aeruginosa* has numerous virulence factors associated with establishing corneal infections(8, 9). These include pathogen-associated molecular pattern (PAMP) such as lipopolysaccharide and flagellin(10, 11), a variety of proteases including elastases (LasA and LasB) and *Pseudomonas aeruginosa* Small Protease (PASP)(8, 12), and the type III secretion system that are important for virulence in experimental models(8, 9, 13, 14). These virulence factors are highly regulated through multiple transcriptional regulators including the LasR quorum sensing master regulator(15). quorum sensing will mediate population density dependent collective responses(16). The most studied among these is the LasR transcription factor. LasR responds to quorum sensing molecule N-(3-oxododecanoyl) homoserine lactone, controls a large portion of the *P. aeruginosa* genome, and is an important regulator of pathogenesis in lung and burn infection models(17-19), as it positively regulates a number of pro-virulence factors including elastase proteases LasA and LasB, and rhamnolipids,(16, 20).

A prior study on the keratitis isolates of *P. aeruginosa* from the Steroids for Corneal Ulcers Trial (SCUT) (21) reported a colony iridescent sheen positive phenotype in 22 of the 101 isolates taken during the course of the study from India(21). This sheen phenotype correlated with significantly worse visual outcomes. These included significantly reduced visual acuity and infiltrate/scar size for patients infected sheen isolates compared to typical *P. aeruginosa*(21). The basis for the sheen phenotype has been shown to be due to mutation of the *lasR* transcription factor gene (22). LasR is a positive regulator of the gene *pqsH*, which codes for an enzyme that converts 2-heptyl-4-quinolone (HHQ) to heptyl-3-hydroxy-4(1H)-quinone (PQS)(22). In the absence of LasR function, HHQ builds up in the cell and creates the sheen phenotype. PQS is an important signaling molecule known as *Pseudomonas* quinolone signal(23). Surprisingly only two nonsynonymous mutations in the *lasR* gene were detected in 21 out of 22 sequenced sheen positive isolates suggesting that the mutations were already present in strains endemic to the country (India). By contrast chronic lung infections, such as those associated with cystic fibrosis, *P. aeruginosa* are frequently observed to gain mutations in *lasR* (22, 24). In the airway, patients are thought to be initially infected by wild-type (WT) *P. aeruginosa*, and *lasR* mutants can then increase over time(25). Like keratitis patients, cystic fibrosis patients infected with *lasR* mutants have been recorded to experience worse disease progression compared to patients infected by wild-type *P. aeruginosa* (26). Together these prior studies suggest that sheen isolates are associated with worse clinical outcomes.

In the SCUT study, all of the sheen isolates tested were isolated between 2006-2010 and caused largely by isolates with one of two *lasR* mutations(21). This study sought to determine whether sheen isolates were a general phenomenon among *P. aeruginosa* keratitis isolates or a geographically isolated observation and if wide-spread whether the same mutant alleles of *lasR* were present and associated with worse visual outcomes. Here we found a concerning increase in *lasR* mutants among the keratitis isolates taken in a tertiary care hospital in the Eastern United States. Data suggests that the mutations were highly variable with one exception, being an insertion element present in several strains; moreover, retrospective analysis suggests that patients with *lasR* mutations had better visual outcomes contrary to the former study.

## Materials and Methods

### Microbiology

De-identified *P. aeruginosa* strains isolated from the corneas of patients with keratitis were retrieved from a clinical tissue bank which is used for validation of diagnostic testing and antibiotic evaluation. The *P. aeruginosa* isolates were collected from 1993 through 2021 by The Charles T. Campbell Ophthalmic Microbiology Laboratory at the University of Pittsburgh School of Medicine and stored at -80°C. *P. aeruginosa* isolates were plated on tryptic soy agar with 5% sheep’s red blood cells and incubated for 18-20 hours at 37°C and sheen phenotype was established visually.

### Molecular Biology

The *lasR* gene was deleted from strain PaC (27) using allelic exchange with plasmid pMQ30, as previously described (28). The plasmid was modified with a Δ*lasR* allele cloned from PA14 Δ*lasR* (21) to generate pMQ767. Primers to amplify the *lasR* region (approximately 500 bp upstream and downstream of the open reading frame) were 4835 and 4836 and listed in Table 1. The resultant strain was verified by PCR and whole genome sequencing. The *lasR* open reading frame was amplified by PCR and cloned into shuttle vector pMQ132 under control of the *Escherichia coli lac* promoter using yeast homologous recombination as previously described (29). Primers to amplify the *lasR* open reading frame were 3217 and 3218 (Table 1). Plasmids were sequenced at the University of Pittsburgh Genomics Core or PlasmidSaurus.

**Table 1.**
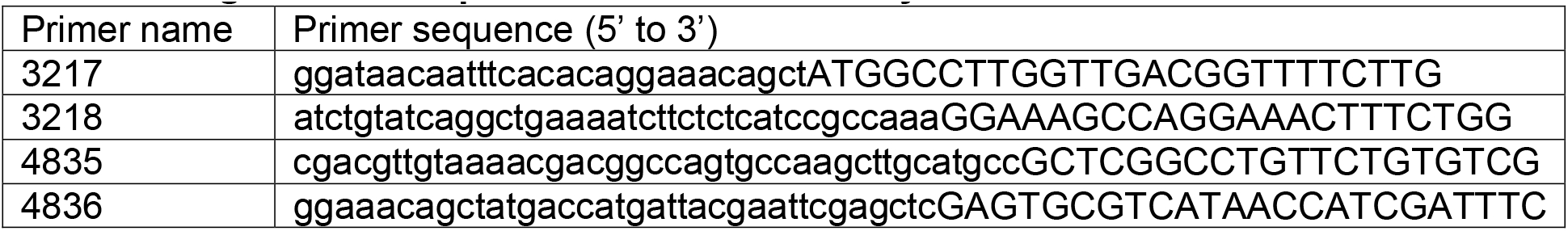
Oligonucleotide primers used in this study.

### Protease Assays

Milk agar plates were made with brain heart infusion and skim milk as previously described (30). Bacteria were incubated at 37°C for 24 hours and zones of clearing were measured on at least three separate occasions. For more quantitative analysis azocasein was used as previously described (31).

### Antibiotic Susceptibility Testing

The minimum inhibitory concentrations (MICs) of *Pseudomonas aeruginosa* keratitis isolates were determined to ciprofloxacin (CIP), tobramycin (TOB), and ceftazidime (CAZ) using E-tests (Fisher Scientific, LIOFILCHEM, MA) on Mueller-Hinton agar as previously described (32). The keratitis isolates tested, taken from the cornea, were chosen arbitrarily out of the deidentified strain bank. The isolates in question were collected anonymously from 2013 to 2022 by the Charles T. Campbell Ophthalmic Microbiology Laboratory and stored at -80°C. The antibiotic susceptibility was determined by comparing the MIC of each to the Clinical and Laboratory Standards Institute breakpoints (33).

### Chart Review

Retrospective review of medical records of all patients diagnosed with culture-positive *P. aeruginosa* keratitis at the University of Pittsburgh Medical Center between 2017 and 2021 was performed. A retrospective review was performed on the medical records of all patients diagnosed with culture-positive *Pseudomonas aeruginosa* keratitis. The study was approved by the Institutional Review Board of the University of Pittsburgh and followed the tenets of the Declaration of Helsinki. Clinical data were collected for each patient, including clinical features, treatment, and outcomes. Demographic features were recorded, including gender and age. Visual acuity was recorded at presentation and after resolution. The visual outcomes were BCVA on the Snellen chart and converted to LogMAR.

### Statistical Analysis

Graph-pad Prism was used to perform Mann-Whitney and ANOVA analysis with Tukey’s post-test, chi-square, and Fisher’s exact tests. P<0.05 was considered significant.

## Results

### An increase in sheen positive *P. aeruginosa* was observed among keratitis isolates from a North Eastern United States tertiary care facility between 1993-2021

*P. aeruginosa* keratitis isolates between 1993-2021 were evaluated for sheen phenotypes on blood agar plates (Figure 1A). The isolate collection and storage approach remained consistent over that time period. A notable increase in sheen positive isolates was observed over the time frame going from 0% between 1993-1997 to 26.2% between 2018-2021 (Figure 1B). All time frames tested were significantly different from 1993-1997 by Fisher’s Exact and chi-square Test and (p<0.01).

**Figure 1.**
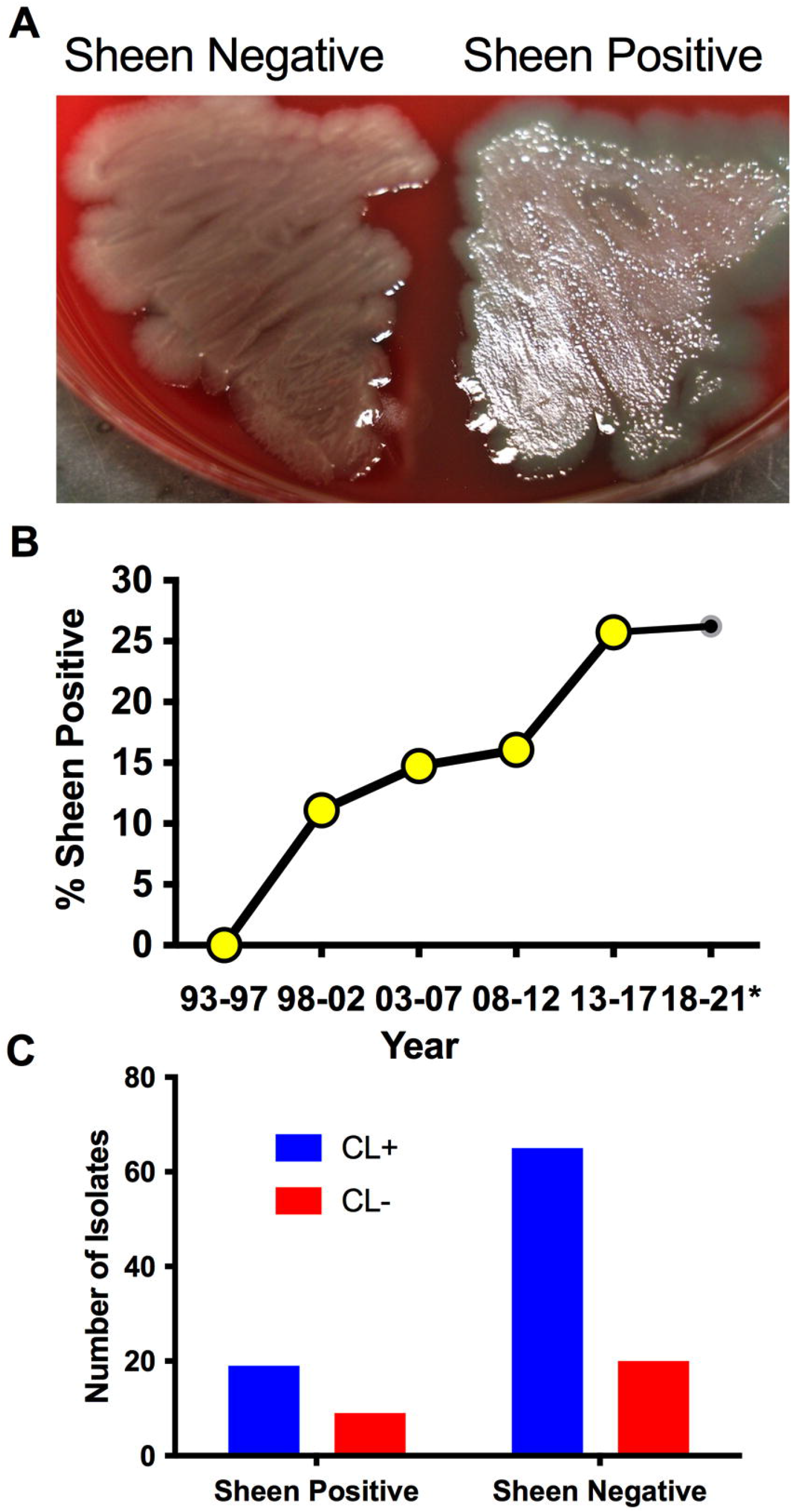
*P. aeruginosa* keratitis isolates with a sheen phenotype are increasing over the last two decades. **A**. Appearance of sheen negative and positive *P. aeruginosa* keratitis isolates on blood agar. **B**. Frequency of PA keratitis isolates with sheen positive phenotype. *, 5 year periods are shown except for 2018-2021. n=399, n≥50 per time period. **C**. Correlation between contact lens (CL) use and sheen status. p=0.455.

Where possible, the contact lens use status of the patient was correlated with the sheen phenotype and no significant difference was observed (p=0.45 by Fisher’s Exact Test) with 23% sheen positive isolates from contact lens associated keratitis (19/84) and 31% sheen positive isolates from non-contact lens associated keratitis (9/29).

We also screened a small library of fluoroquinolone resistant isolates from the New York City area obtained before 2001 (27) and found that two out of six where sheen positive (16.7%), further suggesting that these sheen positive keratitis isolates are a general rather than geographically isolated phenomenon.

### Sequence analysis indicates a variety of mutations and was suggestive of one endemic strain

The *lasR* allele from a subset of arbitrarily chosen strains were cloned into a shuttle vector and sequenced. The prior study with patients from India reported multiple independent isolates with two specific LasR variants I215S (14/22) and P117L (7/22), and one having a missense mutation yielding V221L (1/22) (21). By contrast, our study did not find these mutations and found a wider variety of alterations in the *lasR* sequence. We cloned and sequenced the *lasR* open reading frame (ORF) from 27 strains including 4 from sheen negative isolates and 23 from sheen positive isolates (Table 2). Of the sheen positive strains, 1 had a wild-type *lasR* sequence, others had amino acid substitutions, deletions of insertions that created out of frame mutations, premature stop codons, and insertion elements. Strikingly, 4 of 6 with insertion elements had identical insertions at base pair 126 despite being found over 20 years.

**Table 2.**
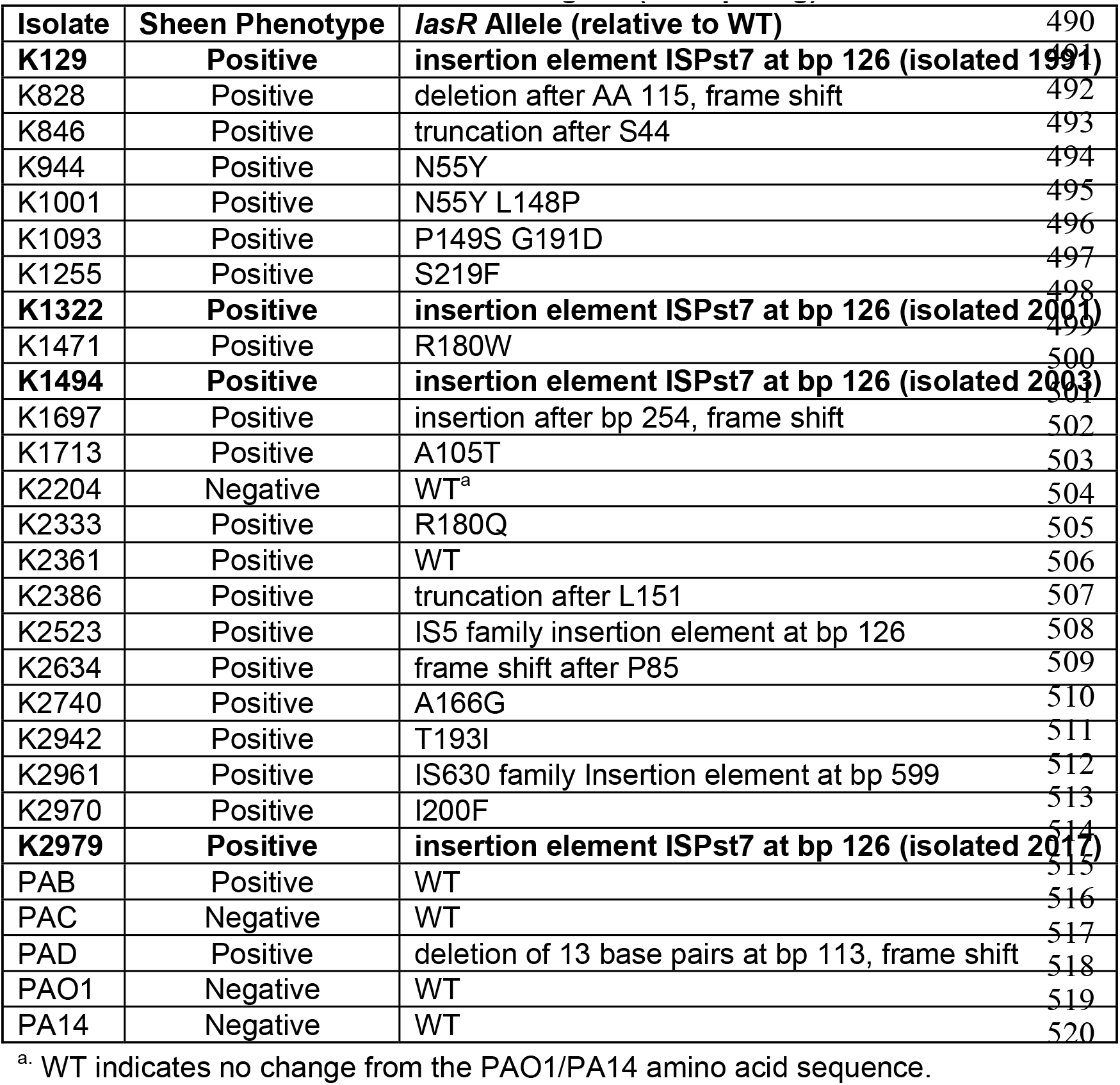
Sequence analysis of PA *lasR* gene (717 bp long).

### Sheen positive keratitis isolates had increased susceptibility rates to a ceftazidime

It has been reported that *lasR* isolates have altered susceptibility to a variety of antibiotics (34-36). To test whether sheen status had an impact on susceptibility to the major topical antibiotics for *Pseudomonas* keratitis (37), we evaluated minimum inhibitory concentrations from keratitis isolates (Table 3). Drugs from three different antibiotic classes were evaluated. There was no difference in the percent susceptible to the fluoroquinolone ciprofloxacin for sheen positive versus negative isolates (92.5% compared to 93.5% respectively, p=0.29 Fisher’s Exact test). For the aminoglycoside tobramycin, there was a non-significant trend toward a higher percentage of susceptible isolates in the sheen negative group which was 10% higher than the sheen positive group (p=0.23). While the cephalosporin ceftazidime is not used as commonly for keratitis, it has been used successfully used to treat keratitis and has been suggested as an alternative for treatment of aminoglycoside and fluoroquinolone resistant isolates (6, 38, 39). For ceftazidime, the sheen negative isolates had a higher frequency of susceptibility than the sheen positive isolates (98.1% vs 85.3% respectively, p=0.01).

**Table 3.** Descriptive statistics of minimum inhibictornycentrations (MICs) for *Pseudomonas aeruginosa* keratitis isolates to ceftazadime, ciprofloxacin, and tobramycin

|  | N | Median (µg/ml) | Mode (µg/ml) | MIC50 (µg/ml) | MIC90 (µg/ml) | Range | Susceptibility (%) |
| --- | --- | --- | --- | --- | --- | --- | --- |
| **Sheen +** |  |  |  |  |  |  |  |
| ceftazidime | 34 | 2 | 1.5 | 2 | 8 | 0.75 - >256 | 85.3 |
| ciprofloxacin | 41 | 0.19 | 0.19 | 0.19 | 0.5 | 0.064 - 1 | 92.5 |
| tobramycin | 41 | 1.5 | 1.5 | 1.5 | 3 | 0.064 - 8 | 20.0 |
| **Sheen -** |  |  |  |  |  |  |  |
| ceftazidime | 103 | 2 | 1.5 | 2 | 8 | 0.75 – 16 | 98.1* |
| ciprofloxacin | 123 | 0.19 | 0.19 | 0.19 | 0.5 | 0.047 – 1.5 | 93.5 |
| tobramycin | 123 | 1.5 | 1.5 | 1.5 | 3 | 0.047 – 5 | 30.6 |

\* significant difference between Sheen + and Sheen – strains by chi-square, p=0.01.

### Sheen positive keratitis isolates are protease deficient

LasR is a known regulator of elastase B (*lasB*) and other proteases thought to aid the bacteria in microbial keratitis(40). The protease activity of arbitrarily chosen keratitis isolates (14 sheen positive and 60 sheen negative) was assessed by measuring the zone of clearance on milk agar plates (Figure 2A). The sheen negative isolates had a zone of 5.6 mm versus 2.7 mm for the sheen positive isolate (p<0.0001, Mann-Whitney test).

**Figure 2.**
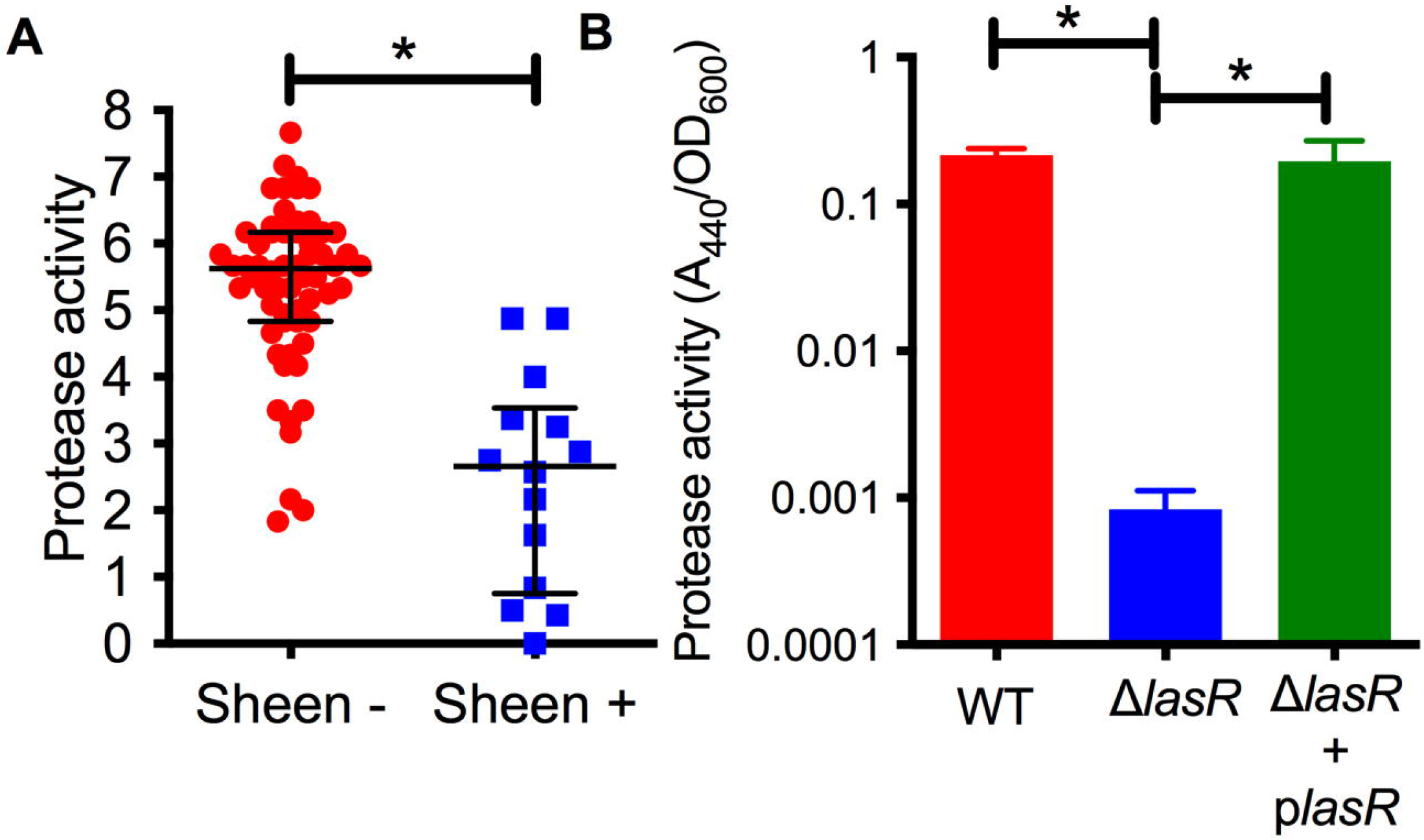
Secreted protease activity is reduced among sheen positive PA keratitis isolates and can be complemented. **A**. Secreted protease activity by PA keratitis isolates. The zone of clearance (mm) on milk plate assay is shown. Each data point indicates the mean zone of clearance for an individual isolate. Medians and IQ ranges are shown. *, p<0.001 by Mann-Whitney. **B**. Secreted protease activity measured by azocasein and normalized by bacterial density from sterile culture filtrates from the clinical isolate PAC (WT) and isogenic Δ*lasR* mutant and the mutant with wild-type *lasR* on a plasmid. The bacteria were grown in LB medium and were harvested at OD_600_=2. Asterisks indicate p<0.01 between indicated groups by ANOVA with Tukey’s post-test, n=3, mean and SD are shown.

To test this further, we generated a *lasR* deletion mutation in strain PaC (27) which is a fluoroquinolone resistant keratitis isolate. The PaC Δ*lasR* mutant was more than 100-fold reduced in protease activity compared to the wild type as measured using azocasein and this defect could be complemented by adding the wild-type *lasR* gene back on a plasmid (Figure 2B).

### The *lasR* alleles from sheen positive isolates were largely loss of function alleles

While the sheen phenotype is linked to *lasR* loss of function mutations, the sheen phenotype can occur due to other mutations such as mutation of the *pqsH* gene which converts HHQ to PQS (41, 42). Several of the cloned *lasR* mutants were tested for function by expression in the PaC Δ*lasR* mutant followed by protease evaluation as a surrogate for LasR function (Figure 3A). Wild-type alleles from PA14 and K2361 restored wild-type levels of protease activity, whereas the *lasR* alleles cloned from sheen positive strains had highly reduced protease activity (Figure 3A).

**Figure 3.**
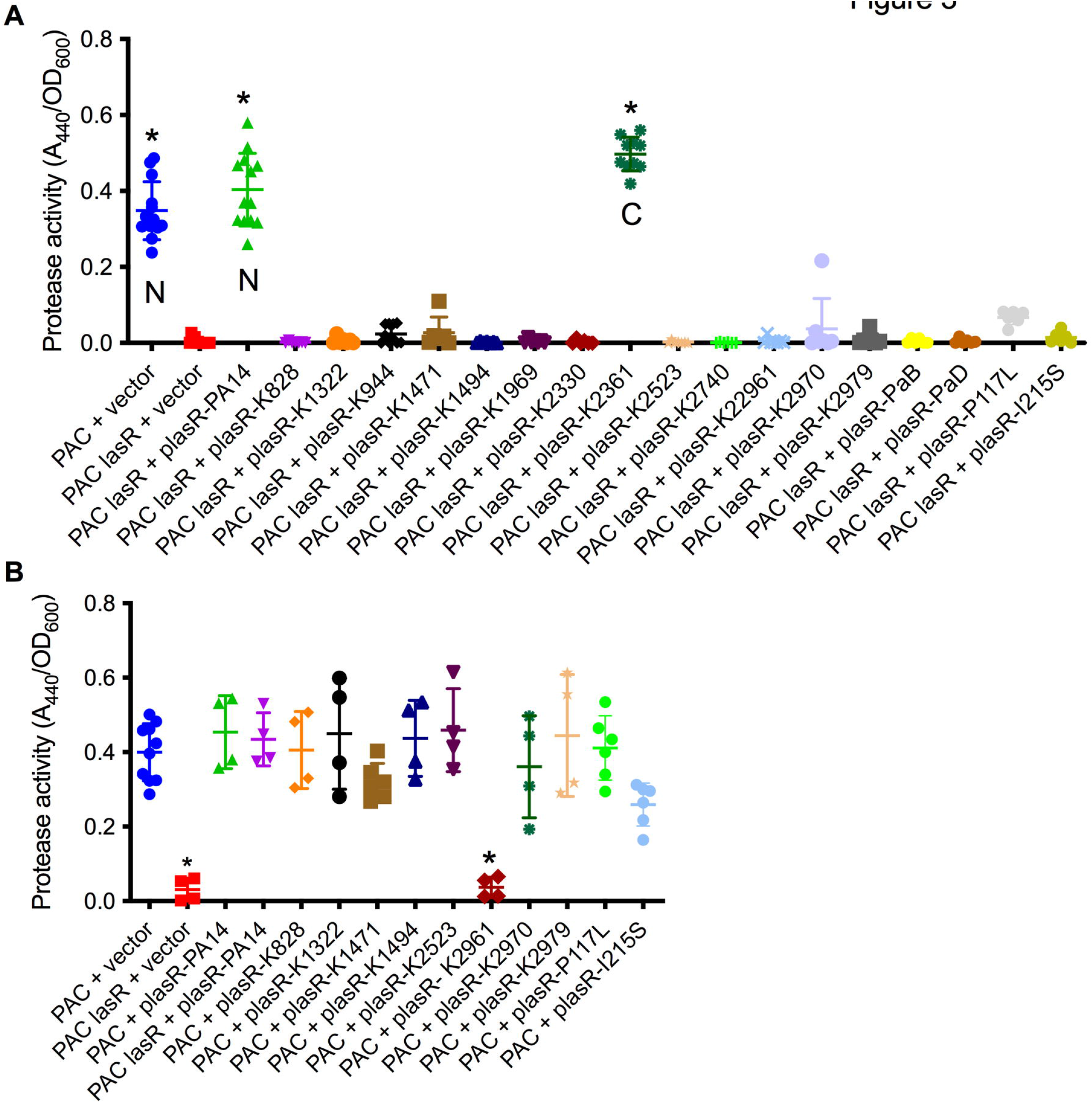
Tested *lasR* alleles from sheen positive isolates were loss of function and generally not dominant negative. **A-B**. Protease activity in supernatants from overnight cultures (18-20h) grown in LB medium was measured using azocasein and normalized by culture density. n≥6, median and standard deviation is shown. **A**. The vector alone negative control (pMQ132), wild-type *lasR* plasmid (plasR-PA14) positive control, and candidate plasmids were expressed in the wild-type PaC to establish base level or PaC Δ*lasR* to determine *lasR* function. “N” indicates a *lasR* allele from a sheen negative strain. “C” indicates *lasR* from a sheen positive isolate with no amino acid changes in the ORF transcript. Asterisk indicates p<0.001 by ANOVA with Tukey’s post-test. **B**. As A, but plasmids were tested in the wild-type PaC to detect dominant negative activity. Asterisk indicates difference from PaC + vector by ANOVA, p<0.01.

Genes for the major two LasR variants I215S (14/22) and P117L (7/22) from the SCUT isolates were also cloned as above, moved into the PaC Δ*lasR* mutant and based on protease activity, both were loss of function mutations (Figure 3A). The PI117L mutant appeared to maintain some activity maintaining 16.6% of wild-type protease activity and 17-fold higher than the vector alone negative control, but was not significantly different than the vector alone negative control. The I215S allele allowed only 3.6-fold higher protease levels than the vector alone negative control.

The dominant negative status of the *lasR* alleles was also tested by expression in the wild-type PaC strain (Figure 3B). Notably, the K2961 allele strongly reduced protease activity in the PaC wild type, whereas the other tested *lasR* alleles did not significantly alter protease activity. The PaC strain with the *lasR*_K2961 allele expressed on a plasmid was remade to ensure that the effect was not artifactual and the reduced protease phenotype was again observed. A similar, although less severe reduction in protease activity was measured when the *lasR*_K2961 allele was expressed in the PA14 wild-type strain (Figure 3B). When the previously described *lasR* alleles from the India isolates were expressed at multicopy in the wild type bacteria, the P117L allele had no effect, but the I215S may have a dominant negative effect with a 43% reduction compared to the wild type expressing the PA14 *lasR* allele (Figure 3B). This difference was not significant by ANOVA with Tukey’s post-test, but a pairwise comparison by Mann-Whitney indicated significance (p=0.0095).

### Clinical outcomes from keratitis isolates with and without sheen phenotype

Where possible the clinical outcomes of patients were determined from clinical records with n=49 for sheen negative isolates and n=19 for sheen positive. Of all of the evaluated variables only final average vision was significantly different (p=0.0106) (Table 4). Surprisingly, unlike the prior study evaluating *lasR* mutant associated keratitis (21), the average visual outcomes were favorable for patients infected with the sheen positive isolates. Though not significantly different, the most severe outcomes, corneal transplants and enucleation, were absent in the sheen positive infected eyes, further suggesting reduced virulence by the sheen positive isolates.

**Table 4.**
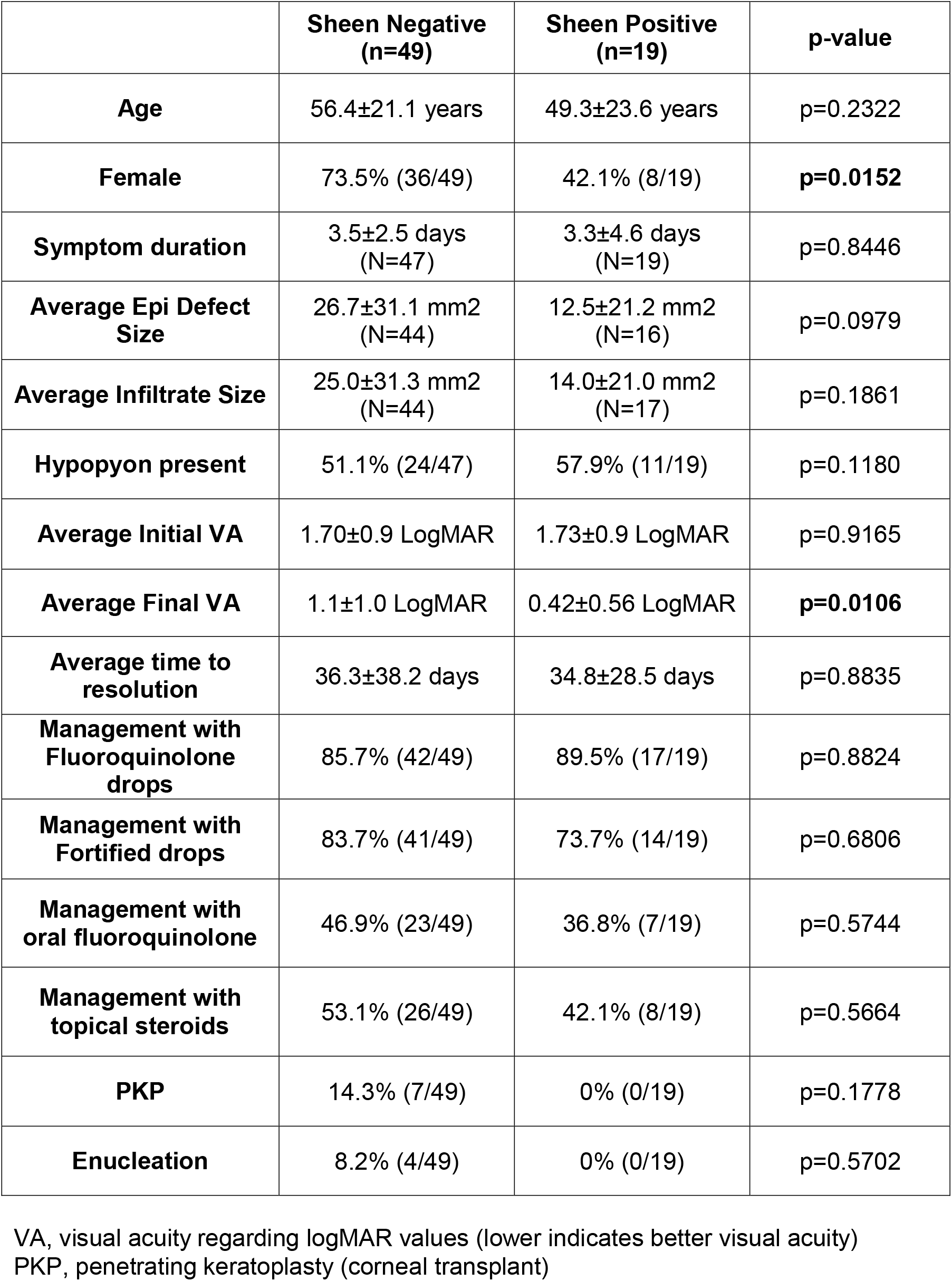
Clinical outcomes from keratitis patients.

## Discussion

This study has demonstrated an increase in sheen positive *P. aeruginosa* keratitis isolates in a tertiary care hospital in the Eastern United States. This suggests that the abundance of sheen positive isolates is a general rather than a regional phenomenon. The reason for the increase observed in this study was not clear. The clinical microbiologist that collected the samples maintained the same collection protocol over the period of isolate collection, so differences in this would not account for the increase in sheen isolates.

Another consideration was whether the sheen positive isolates from our study are *lasR* mutants. The majority (22 out of 24) of sheen isolates that were sequenced had changes in the *lasR* sequence and those tested did not code for functional proteins. Because only the open reading frames were cloned, other mutations that render the strain LasR-deficient could be missed, for example, promoter mutations. However, it is formally possible that in a small subset of the keratitis isolates, LasR-independent changes could be responsible for a subset of the sheen isolates. Therefore, we conclude that the majority if not all sheen positive keratitis isolates have defects in LasR function.

The Hammond study indicated worse visual outcomes for patients with sheen positive isolates (21), by contrast patients in this study had strikingly better visual acuity, as well as no incidence of the severe outcomes of enucleation and corneal transplantation that were present in the sheen negative infected patients. The reason for this discrepancy is not clear, but could possibly be due to the different strains or the specific mutations associated with strains isolated in the SCUT study. Moreover, the SCUT study used a standard protocol for timing and methodology for obtaining visual acuity measurements, which lends more weight to that analysis.

Nevertheless, the reduced severity observed in the clinical data from this study were consistent with a recent paper using a rabbit corneal infection model that demonstrated reduced corneal perforation and bacterial proliferation of a *lasR* deletion mutant of strain PA14 compared to an isogenic wild type suggesting that LasR promotes keratitis severity (43). Studies with mice show mixed results with strain PA01 with *lasR* deletion mutations. The Pier group reported that C3H/HeN mice with scarified corneas required fewer *lasR* mutant bacteria to cause keratitis compared to the wild type (44). Whereas the Willcox group used the same bacterial strains with BALB/c mice and found indistinguishable infection frequencies for both bacterial strains, but reported that bacterial proliferation and severity scores were reduced in eyes infected with the *lasR* mutant (44).

In conclusion, *lasR* mutant *P. aeruginosa* appear to be increasing among keratitis patients and this may be a world-wide phenomenon. The highly variable nature of the *lasR* mutations among isolates in our study does not indicate whether the strains mutated during infection or prior to infection in general. The exception being identification of multiple isolates with a mutation at base pair 126 which suggests the existence of a regional endemic strain. Interestingly, the identical insertion element was reported in the *lasR* gene from a *P. aeruginosa* isolated from a bean plant in Spain suggesting an environmental source (45). Whether these strains are more or less virulent is in question and more research is needed to determine the level of concern, however, our study suggests that the sheen positive *lasR* isolates are less pathogenic.

## Author Contributions

RMQS – writing, conceptualization, methodology, funding; SA – writing, methodology; NA – methodology, review and editing; CVS - methodology, review and editing; AG - methodology, review and editing; HQS - methodology, review and editing; SMM - methodology, review and editing; DKD - conceptualization, review and editing; AM - conceptualization, review and editing; JDC - methodology, review and editing; RCC - methodology, review and editing; EGR - conceptualization, review and editing; RPK – conceptualization, methodology, review and editing; MEZ - conceptualization, writing, review and editing; VJ - conceptualization, writing, review and editing.

## Acknowledgments

The authors thank Deborah Hogan at Dartmouth Medical School for strains.

## Data Availability Statement

The original information presented in this study are included in the article and further inquiries can be directed to the corresponding author.

